# Characterisation of a type II functionally-deficient variant of alpha-1-antitrypsin discovered in the general population

**DOI:** 10.1101/452375

**Authors:** Mattia Laffranchi, Emma L. K. Elliston, Fabrizio Gangemi, Romina Berardelli, David A. Lomas, James A. Irving, Annamaria Fra

## Abstract

Lung disease in alpha-1-antitrypsin deficiency (AATD) results from dysregulated proteolytic activity, mainly by neutrophil elastase (HNE), in the lung parenchyma. This is the result of a substantial reduction of circulating alpha-1-antitrypsin (AAT) and the presence in the plasma of inactive polymers of AAT. Moreover, some AAT mutants have reduced intrinsic activity toward HNE, as demonstrated for the common Z mutant, as well as for other rarer variants. Here we report the identification and characterisation of the novel AAT reactive centre loop variant Gly349Arg (p.G373R) present in the ExAC database. This AAT variant is secreted at normal levels in cellular models of AATD but shows a severe reduction in anti-HNE activity. Biochemical and molecular dynamics studies suggest it exhibits unfavourable RCL presentation to cognate proteases and compromised insertion of the RCL into β-sheet A. Identification of a fully dysfunctional AAT mutant that does not show a secretory defect underlines the importance of accurate genotyping of patients with pulmonary AATD manifestations regardless of the presence of normal levels of AAT in the circulation. This subtype of disease is reminiscent of dysfunctional phenotypes in antithrombin and C1-inibitor deficiencies so, accordingly, we classify this variant as the first pure functionally-deficient (type II) AATD mutant.

## Background

Severe alpha-1-antitrypsin deficiency (AATD, MIM #613490) affects approximately 1 in 2000 of the Northern European population. It is associated with pathogenic variants of the *SERPINA1* gene (MIM #107400) which encodes alpha-1-antitrypsin (AAT). AAT is the archetypal member of the serpin superfamily of serine-protease inhibitors [1]; its primary physiological role is to protect the lung parenchyma from attack by the serine proteases neutrophil elastase (HNE), cathepsin G [2] and proteinase 3 [3]. Mutations that reduce AAT plasma levels alter the balance between inhibitory and proteolytic activity leading to early onset emphysema and COPD [4]. A subset of *SERPINA1* pathogenic alleles, well represented by the common severe deficiency Z allele (E342K, p.E366K) can lead to accumulation of the protein as ordered polymers within the endoplasmic reticulum (ER) of hepatocytes [5], predisposing ZZ homozygotes to liver disease [6]. AAT polymers have been also found in the circulation of AATD patients with different genotypes [7,8]; they are known to exert pro-inflammatory functions by stimulating neutrophils and monocytes [9] and their presence is likely to over-estimate the amount of active AAT in the circulation.

The protease inhibition mechanism has been extensively studied in AAT as well as in other serpins. A specialized region of the serpin structure, termed the reactive centre loop (RCL), acts as pseudo-substrate for a cognate protease. The critical amino acids for the “bait” sequence of the AAT RCL are the P1-P1’ residues M358 and S359 (according to the nomenclature in [10]). After docking of the protease to the P1 residue in the RCL of the AAT molecule, the protease cleaves the P1-P1’ peptide bond, forming an acyl-intermediate bond with the backbone carbon of the P1 residue. The cleavage is followed by a re-arrangement of AAT from a “stressed” to a “relaxed” structure, which flips the protease from the upper to the lower pole of the serpin as the RCL inserts as an extra strand in β-sheet A. Following auto-insertion of the RCL, the catalytic triad of the enzyme becomes distorted, leading to its irreversible inactivation [11]. This complex is cleared from the circulation the liver with a mechanism of re-uptake mediated by members of the lipoprotein receptor family [12].

Extensive biochemical and structural analyses of the inhibitory mechanism [13–18] have provided evidence for multiple intermediate states [15]: an initial docked “Michaelis” complex with the protease, cleavage of the reactive centre loop, insertion of the loop into the breach region at the top of β-sheet A, perturbation of the F-helix to allow passage of the protease, completion of loop insertion and finally compression of the protease. Interference at any of these steps can affect the inhibitory process by altering the balance between productive complex formation and non-productive cleavage of the serpin and release of the protease (Fig 1) [19]. Some AAT mutants manifest an intrinsic reduced anti-protease activity. The common Z variant, whose mutation is located in the breach region, shows an impaired activity compared to wild-type M AAT [20]. Other rarer AATD-associated variants also reduce the inhibitory efficiency for HNE, with an increased non-productive turnover of inhibitor for Queen’s (p.K178N, K154N) and Baghdad (p.A360P, A336P) mutants and decreased rate of interaction seen with F AAT (p.R247C, R223C) [21–24]. In contrast, the AAT variant Pittsburgh (p.M382R, M358R) contains a substitution at P1 that switches specificity from the inhibition of HNE to inhibition of the clotting factors thrombin (THR) and factor XIa (FXI), resulting in fatal bleeding events [25].

**Fig 1.**
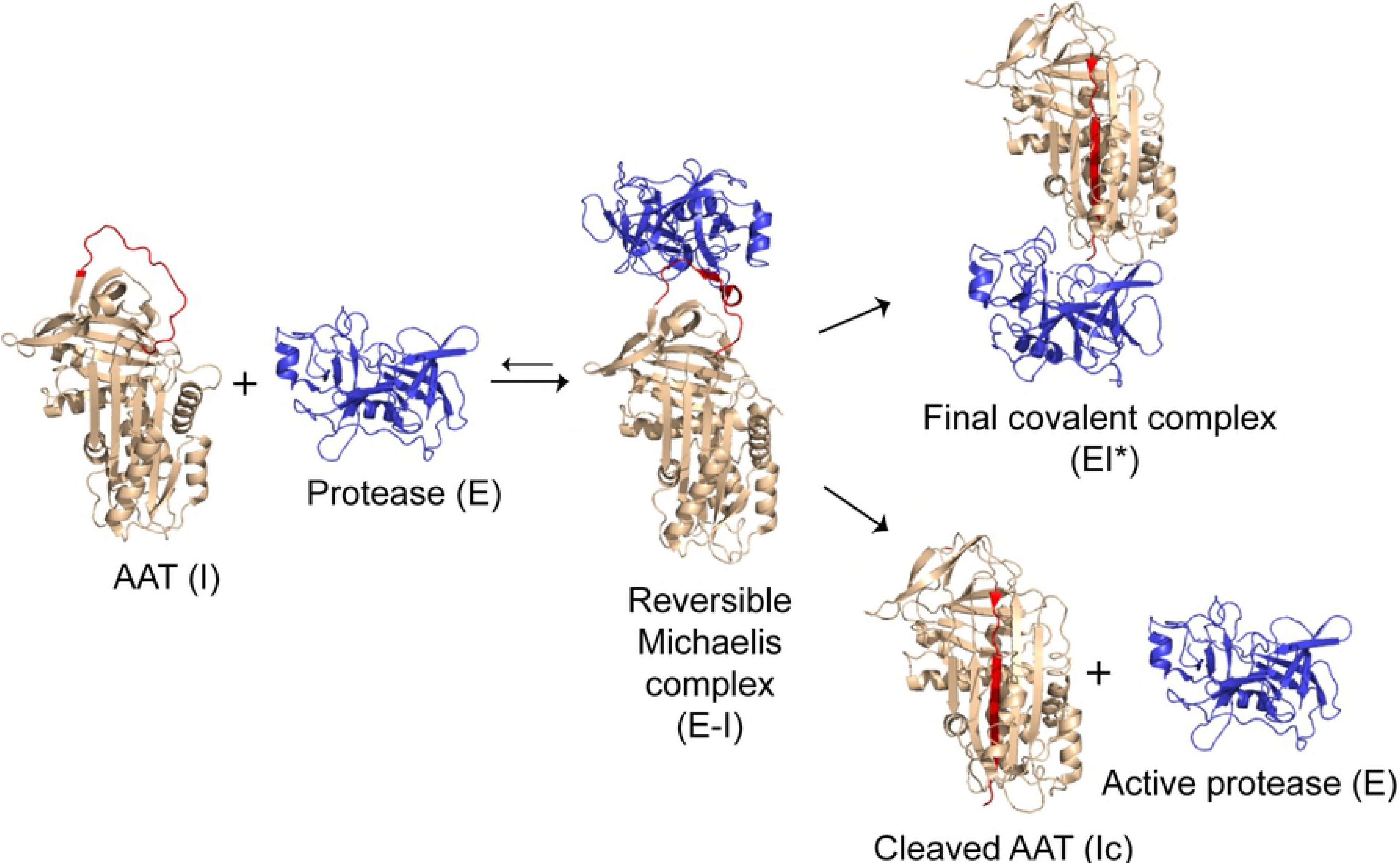
Inhibitory mechanism of alpha-1-antitrypsin. A schematic illustrating the inhibitory mechanism of alpha-1-antitrypsin (AAT), showing the progression from active inhibitor (I, PDB: 1QLP) and active protease (E, PDB: 1OPH), to the formation of the reversible Michaelis complex (E-I, PDB: 1OPH), and the branched pathway that leads to irreversible complex formation (EI*, PDB: 2D26) or cleaved inhibitor (Ic, PDB: 1EZX) and active enzyme (E). The figure was prepared with PyMOL (The PyMOL Molecular Graphics System, Version 2.0 Schrödinger, LLC).

The ExAC consortium [26] is a free-to-access repository presently containing about 60,706 exome sequences, gathered from several human populations. By querying this database, it is possible to identify novel putatively pathogenic variants of proteins including AAT [27]. From this and other human population databases, we have evaluated novel *SERPINA1* variants that fall within the RCL domain - a region critical to the inhibitory mechanism - with the aim of identifying novel dysfunctional variants in the general population.

By this approach, we have identified the Gly349Arg (p.G373R) variant in the RCL region. Using expression in mammalian cells, biochemical and structural analyses we show that this variant is secreted at wild-type levels but lacks anti-protease activity. These data support the classification of Gly349Arg as the first type-2 deficiency variant of AAT to be described.

## Results

### Search for putatively dysfunctional alpha-1-antitrypsin variants in human populations

With the aim of identifying novel dysfunctional *SERPINA1* mutants, we considered *SERPINA1* variants annotated in population databases that fall within the AAT RCL domain (between residues 344 and 362 in the canonical nomenclature). We identified a Gly→Arg mutation at the P10 (349) position that violates the pattern noted for residues in the RCL hinge region [28], and therefore putatively could interfere with inhibitory activity (Fig 2A). In addition, Gly349 is relatively conserved among *SERPINA1* orthologues, while within the human serpins the 349 position shows a preference for small aliphatic residues (Fig 2B and S1 File). There was sufficient information in the ExAC database to establish that this variant was present in three separate carriers of 60,706 individuals, and they were not first- or second-degree relatives. A female and a male were present in the Non-Finnish ExAC population, and a male in the Finnish population. We received age data for two of the three individuals: one 40-45 years old, and another 65-70 years old.

**Fig 2.**
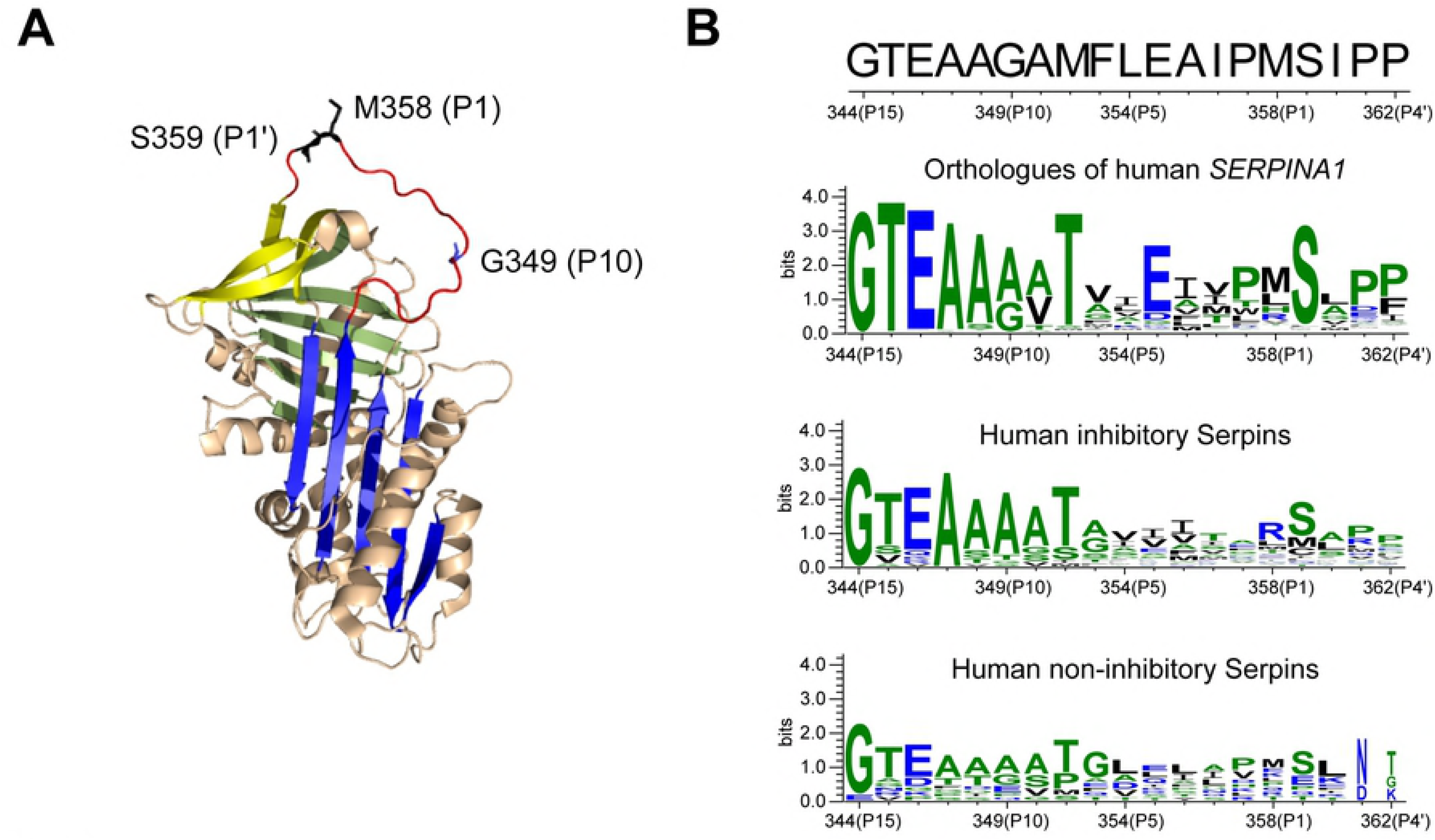
Glycine 349 localization and conservation within the alpha-1-antitrypsin reactive centre loop. **(A)** The Met358 (P1) and Ser359 (P1’) residues, critical for the anti-protease activity of AAT, are shown as black sticks on the native AAT structure (PDB: 1QLP), while the residue Gly349 (P10) is showed as blue stick. β-sheet A is blue, β-sheet B is green and β-sheet C is yellow; the RCL is coloured in red. The figure was prepared with PyMOL. **(B)** Conservation of residues in the RCL (top sequence, from residue G344 to P362) is represented using WebLogo [56], calculated from a sequence alignment of the *SERPINA1* orthologues or from human serpins paralogues.

### Characterization of the G349R alpha-1-antitrypsin variant in cell models

Several cellular models have been utilized successfully to study accumulation-prone variants of AAT and other serpins, and have been shown to represent excellent predictors of *in vivo* behaviour [7,29–35]. To characterize the efficiency of secretion of AAT G349R, we transiently expressed it in HEK293T and Hepa 1-6 cell models in comparison with the wild-type M AAT and the common deficient and polymerogenic Z mutant of AAT (Fig 3).

**Fig 3.**
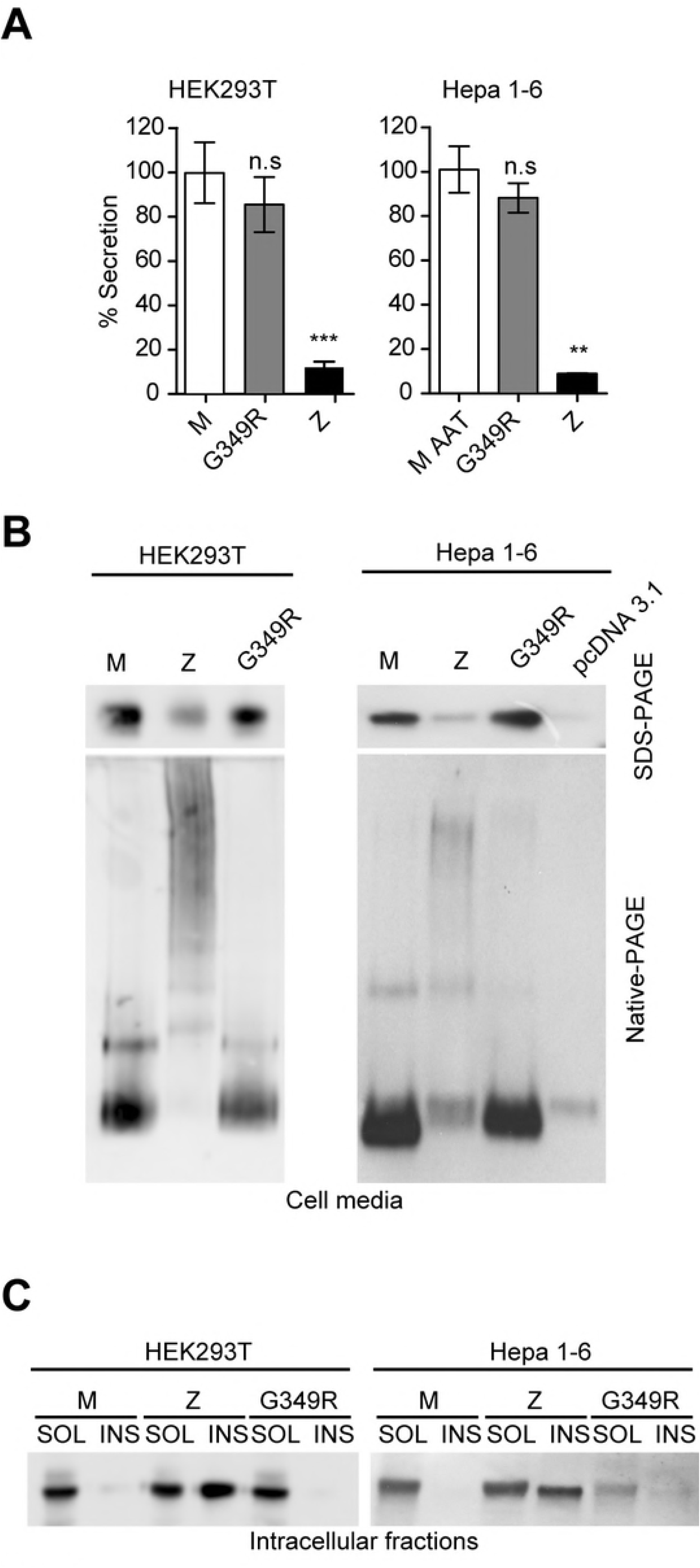
Characterization of the G349R variant in cell models. **(A)** AAT levels in cell media from transfected HEK293T or Hepa 1-6 cells were quantified by sandwich ELISA and represented as percentages of wild-type M levels (mean ± SD, *n* = 3; one-way ANOVA, *p* < 0.0001; two-tailed unpaired t-test between each variant and M AAT, n.s nonstatistical significative, **p < 0.001, ***p < 0.0001). **(B)** Immunoblots with anti-total AAT pAb loaded with equal volume of cell media from HEK293T (left) or Hepa 1-6 (right) cells expressing the indicated variants, resolved by 7.5% w/v acrylamide SDS-PAGE (top) and 8% Native-PAGE (bottom). **(C)** Immunoblots with anti-total AAT pAb of NP40-soluble (SOL) or insoluble (INS) cellular fractions from HEK293T (left) or Hepa 1-6 (right) cells expressing the indicated variants, resolved by 7.5% SDS-PAGE.

AAT levels in the cell media from HEK293T and Hepa 1-6 transfected cells were determined by ELISA (Fig 3A) showing that in both cell lines, AAT G349R appears to be secreted at similar levels of the wild-type M AAT (85.6% ± 12.5 MEAN±SD for HEK293T cells, 87.3 ± 6.6 for Hepa 1-6 cells).

Analysis of the extracellular AAT by non-denaturing PAGE also shows that AAT G349R is exclusively monomeric in the cell media (Fig 3B), while the Z AAT polymerogenic and deficient variant is present in the media primarily as high molecular weight oligomers.

In addition, we investigated the intracellular distribution of AAT between the NP40-soluble and - insoluble fractions. As previously reported [33], the tendency of an AAT variant to accumulate in the NP-40 insoluble fraction is a symptom of severe polymerogenic tendency. In Fig 3C the only AAT variant to precipitate within the insoluble fraction is Z AAT.

In conclusion, the G349R variant does not exhibit a “classical” deficiency phenotype.

### The G349R alpha-1-antitrypsin variant is non-functional

A survey of the literature identified a protein engineering study in which the P10 alanine residue of α_1_-antichymotrypsin was arbitrarily mutated to arginine to evaluate its ability to become incorporated into β-sheet A upon cleavage. It was found that due to compensatory molecular rearrangements of the α_1_-antichymotrypsin molecule, there was an increase in substrate behaviour rather than an elimination of inhibitory activity [36]. Correspondingly, we undertook experiments to assess the functional capability of G349R AAT, with a view to ascertaining the clinical impact of this variant.

One of the characteristics of the serpin-enzyme complex is that it is stable in the presence of SDS [13,27]. We incubated the media of HEK293T cells expressing M or G349R with an equimolar or 2:1 excess over HNE (Fig 4A) and analysed complexes by SDS-PAGE and immunoblot. Instead of forming the 68 kDa SDS-resistant complex as the wild-type M (Fig 3A, top arrow), the G349R variant was preferentially cleaved by the HNE (Fig 4A, bottom arrow).

**Fig 4.**
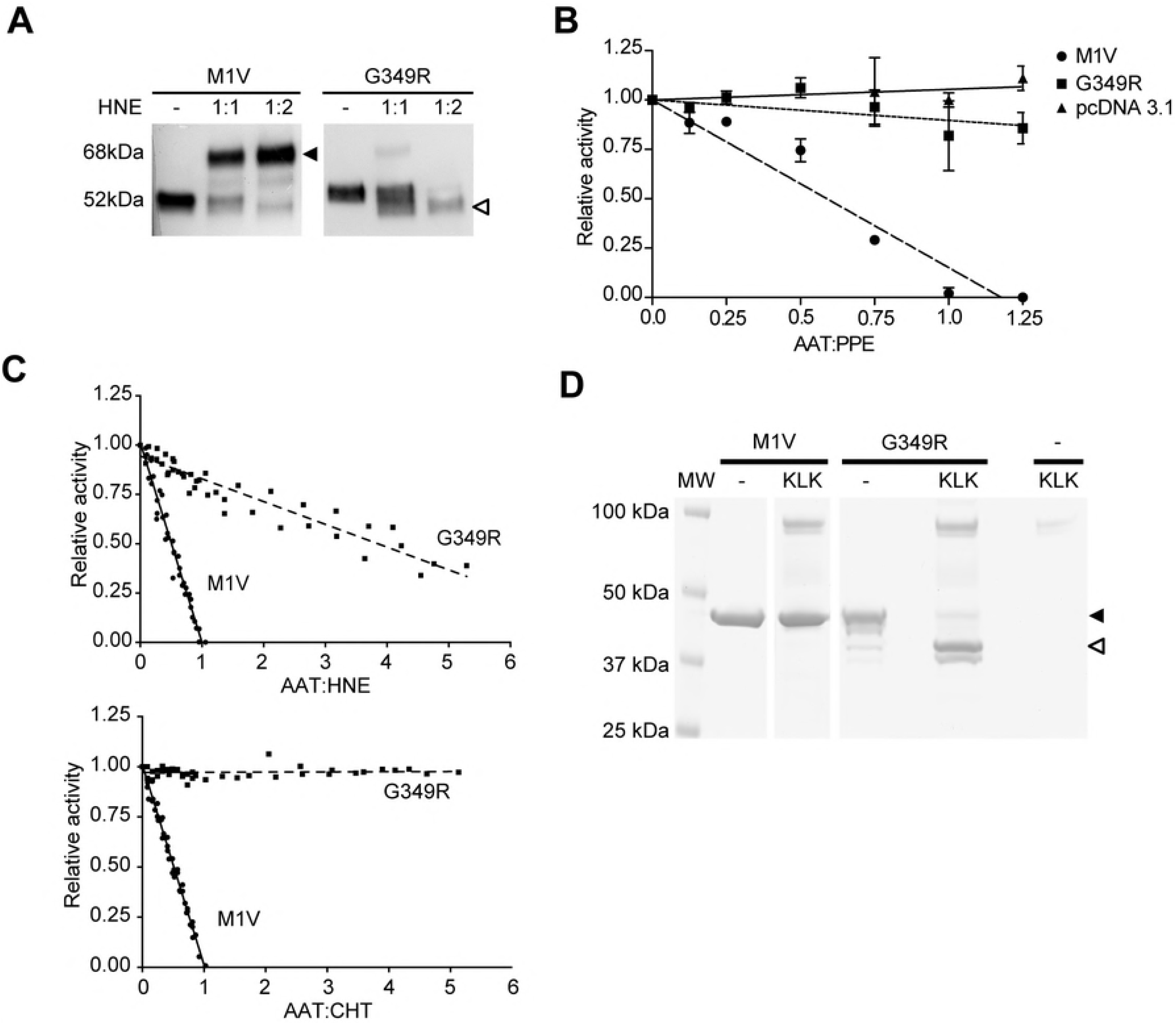
The AAT G349R variant has a defective inhibitory mechanism. **(A)** Cell media of HEK293T transfected cells were incubated with (+) or without (−) equimolar or double concentration of neutrophils elastase (HNE) for 30 min at 37°C, and the complexes (68 kDa, dark arrow) were resolved from unreacted AAT monomers (52 kDa) or cleaved forms (48 kDa, white arrow) by 7.5% w/v acrylamide SDS-PAGE and immunoblot with anti-AAT pAb. **(B)** An enzymatic assay using the pSuccAla3 chromogenic substrate with porcine pancreatic elastase (PPE) that had been preincubated with an increasing ratio of either wild-type M1V (dashed line) or Iners AAT (dotted line) AAT variants from HEK293T cell media. pcDNA (solid line) represents the media of HEK293T cells not expressing AAT (n=2, One-way ANOVA, p-value < 0.017). **(C)** Relative protease activity in the presence of increasing ratio of recombinant AAT wild-type M1V (solid line) or G349R variants (dashed line). In the left panel summarises the experiments using HNE, while the right panel shows the inhibition of chymotrypsin (CHT) (n=2). **(D)** Recombinant AAT wild-type M1V and G349R variants were incubated with (+) or without (−) an equimolar concentration of human purified plasma kallikrein (KLK) for 30 min at 37°C, and the samples were resolved by 4-12% w/v acrylamide SDS-PAGE. Arrows point to the uncleaved (48 kDa, dark arrow) and the cleaved forms (44 kDa, white arrow).

To further confirm this dysfunctional behaviour, we measured the activity against porcine pancreatic elastase (PPE), a tool serine protease sometimes used as a surrogate for HNE. After incubation with an increasing ratio of AAT variants in the media of HEK293T cells, compared to the inhibitory activity of the wild-type M (dashed line), the new variant (dotted line) showed no signs of inhibitory activity (Fig 4B).

The variant was also expressed in bacteria and purified to homogeneity. It was found that this material required an 8-fold greater concentration to fully inhibit HNE than M AAT, representing an around 80% decrease in activity (Fig 4C, left panel), and was entirely inactive against the tool protease chymotrypsin (CHT) (Fig 4C, right panel). The exclusive presence of the cleaved form of G349R AAT when tested with HNE (Fig 4A) shows that the loop is still recognised by the protease, but that the branched inhibitory pathway is skewed in favour of non-productive turnover of AAT and premature release of active protease.

AAT Pittsburgh represents a variant in which an amino acid substitution in the specificity-determining site of the RCL introduces a novel cleavage site for other human proteases, including thrombin [25]. While the G349R mutation does not prevent an interaction with HNE, it may similarly lead to off-target recognition by other proteases. By querying the MEROPS database [37] for human proteases that could potentially cut the sequence introduced by the G349R substitution (EAA**R**AMFL), we identified plasma kallikrein (KLK), which recognises the P1-P1’ sequence R-X, as a candidate. Using the recombinant protein, it was found that indeed this variant is susceptible to cleavage by KLK when incubated at equimolar ratio (Fig 4D).

In conclusion, AAT G349R is a novel dysfunctional variant lacking effective inhibitory activity. Many AAT variants are named according to the birthplace of the proband or the site of the diagnosing centre; as this information is unknown we decided to name the novel variant AAT Iners, from the latin word meaning “inactive”.

### Comparative structural analysis and molecular dynamics simulations suggest an impaired presentation and insertion of the RCL domain

The experimental data suggest two possible structural causes of dysfunction: the first is that the conformation of the RCL is altered, which can affect protease recognition and docking; the second is that the bulky charged group that replaces Gly349 impedes a timely conformational change of the RCL after cleavage.

An Ala→Arg mutation has been introduced at the equivalent site in a protein engineering study of a related serpin, α_1_-antichymotrypsin [36]. A comparison between the crystal structures of the RCL-cleaved forms of Ala349Arg alpha-1-antichymotrypsin (PDB ID: 1AS4) and the wild-type protein (PDB ID: 2ACH) showed that the introduced arginine side-chain can be accommodated into β-sheet A, but only by rearrangement of the local hydrophobic packing interactions and incorporation of a counter-ion [36]. This is sufficient to greatly reduce inhibitory efficiency, as measured by the non-productive turnover of inhibitor [36]. A comparison with the equivalent residues in cleaved AAT (PDB ID: 1EZX) shows that in order to accommodate the G349R mutation, similar shifts would be required in Phe384 and Ala336, and one residue, Phe51, provides a more marked occlusion in AAT with respect to Ile51 in alpha-1-antichymotrypsin (Fig 5A).

**Fig 5.**
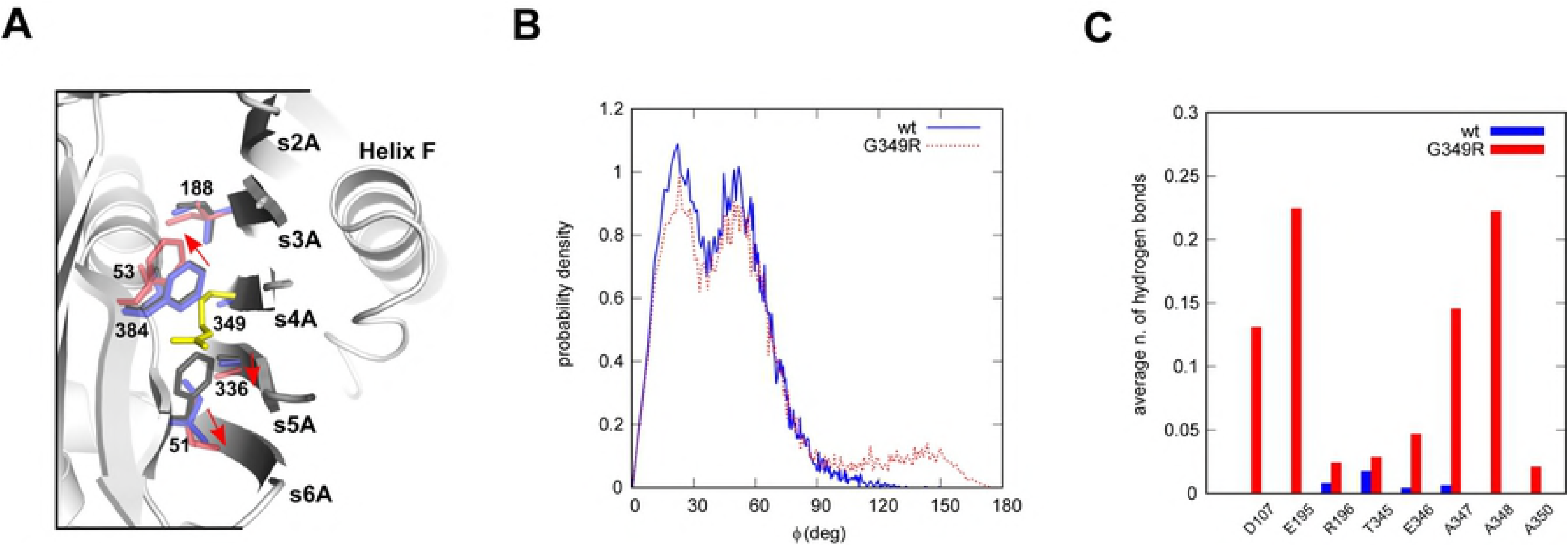
Comparative structural analysis and molecular dynamics simulations suggest an impaired presentation and insertion of the RCL. **(A)** Residues in the vicinity of position 349 (yellow) following loop insertion are shown as sticks for cleaved wild-type alpha-1-antichymotrypsin (PDB: 2ACH, purple), the A349R variant (PDB: 1AS4, red) and AAT (PDB: 1EZX, black). Red arrows show the compensatory side-chain movements that occur in alpha-1-antichymotrypsin to accommodate the bulky charged arginine residue. The figure was prepared using PyMol. **(B)** Probability distribution of the angle φ defined in the text, as calculated in MD simulations for the wild type (solid line) and the G349R variant (dashed line). **(C)** Average number of hydrogen bonds between residue 349 and selected residues calculated in the MD simulations, for the wild type (wt, black) and the AAT Iners variant (G349R, grey).

As well as interference with β-sheet incorporation during inhibition, a glycine-to-arginine mutation would be expected to result in angular restriction of the local protein backbone in the RCL-exposed conformation. To test this possibility, molecular dynamics (MD) simulations were performed for the wild-type protein (PDB ID: 1QLP) and AAT Iners in the uncleaved form. In each case, four 70-ns simulations were performed, and the last 50 ns of each trajectory were combined to form an aggregated 200-ns trajectory for subsequent analysis. Careful visual analysis of the simulations showed that the orientation of the side chain of Met358 in the AAT Iners variant has a different behaviour with respect to the wild-type (S1 Video). To quantify this phenomenon, an observable was defined as follows: a vector **v**_1_ was defined, joining atom Cα of residue Phe370 to atom Cα of residue Met358, and a vector **v**_2_ joining atom Cα to atom S of residue Met358. The angle φ between vectors **v**_1_ and **v**_2_ was then calculated at each time during the simulations. The resulting probability density distribution, in the range from 0° to 180°, is shown in Fig 5B. Values of φ near 0° correspond to the side chain of Met358 pointing outwards with respect to the bulk of the protein, while values near 180° correspond to the side chain pointing inwards. The probability of finding an inward side chain (90° < φ < 180°) was 2% for the wild type and 11% for AAT Iners. To further support the atypical behaviour of the RCL region with the G349R substitution, we performed simulations of the protein immediately following cleavage, and calculated the average number of hydrogen bonds between Arg349 and selected residues (Fig 5C). Arg349 acquired novel interactions with the proximal residues Ala347 and Ala348, but also established additional bonds with the distal amino acids Asp107 and Glu195. Thus, AAT Iners is predicted from the simulations to have an altered RCL presentation. Coupled with the observation that compensatory movements are required during insertion into β-sheet A, possibly including negation of the buried charge, this substitution leads to malfunction of early steps in the inhibitory mechanism.

## Discussion

The reactive centre loop is an exposed loop that exhibits relatively few interactions with the serpin body and plays a lesser role in stability and folding of the native molecule than other functional components of the serpin structure. As the primary determinant of serpin specificity, this element is by definition permissive of sequence variability, with the exception of marked deviations in length [38].

However, inhibitory serpins exist as two conformations and must present a sequence that is recognised by the target protease, and these facts impose evolutionary constraints on some positions in the loop [1,28]. Thus, mutations at certain positions can impair the interaction with a target protease, permit non-productive degradation by other proteases, or perturb the inhibitory mechanism by interfering with the accommodation of the RCL by β-sheet A.

An increasing number of gene variants are being discovered by exome or genome-wide sequencing of large cohorts and made available in annotated databases such as ExAC or LOVD [26,27,39].

In the present study, we have shown that one such variant, G349R, is secreted normally in the native form by cells, but would provide no protection against proteolytic activity due to its near-inability to form a stable inhibitory complex with target proteinases. Notably, this variant was predicted by the pathogenicity predictor REVEL as benign (score 0.448 with pathogenicity threshold at 0.477 [27]), suggesting that mutations affecting the RCL and the activity may be misinterpreted by commonly-used predictors.

Upon cleavage of the P1-P1’ bond by a protease, the RCL of AAT begins its insertion into β-sheet A via a zipper-like motion involving residues 342 to 350 [13]. To achieve the full insertion of the RCL into β-sheet A, the AAT molecule must rearrange helix-F (hF) [14,15], a step predicted by accelerated molecular dynamics to critically involve Ala348 and Gly349 [14].

Ultimately, the RCL is accommodated into the β-sheet and the protease is trapped. Interference with the timely progression of this change is deleterious to inhibitory activity. One member of the serpin superfamily unable to spontaneously undergo the loop-to-sheet conformational change is ovalbumin [40], which has an arginine residue at the P14 position of the RCL. When incorporated into inhibitory serpins, this residue abrogates activity [41]. Conversely, replacement by a serine or threonine in ovalbumin greatly improves the insertion rate of the RCL upon cleavage by a protease [40,41]. Mutations at other positions violating the hinge region motif at the ‘P-even’ residues in AAT have similarly shown compromised inhibitory activity [28,42,43], and studies using its closest homologue, α_1_-antichymotrypsin [1], have shown loss of activity from introduction of arginines at P14, P12, and P10 positions [36]. In all three cases, structure determination by protein crystallography revealed that the RCL could be accommodated, in the case of P14 by introducing a twist in the backbone orienting the side-chain towards the solvent, and in the latter two by compensatory changes in the hydrophobic residues underlying β-sheet A. The structural evidence that such unfavourable substitutions can be accommodated highlights that the partition between a productive and non-productive interaction is contingent on the balance between the kinetics of insertion and hydrolysis [19].

The Pittsburgh AAT variant represents a case in which substitution within the specificity-determining residues results in the ability to interact with a novel target protease. Changes outside of this region can also lead to aberrant interactions but are much more likely to lead to nonproductive cleavage. Our data show that AAT Iners can act as a novel substrate of KLK, which liberates bradykinin (BK), a vasodilator hormone, from the high molecular weight kininogen (HMWK) [44] and activates plasminogen [45]. Given the expected high circulating AAT concentrations for a heterozygote *in vivo* (around 10 to 25μM) this could conceivably lead it to act as a competitive substrate for this enzyme.

Two other genetic disorders have been reported that are associated with dysfunctional plasma inhibitory serpins produced at normal levels. Hereditary angioedema type I and II (HAE, #MIM106100), an autosomal dominant disorder, is caused by mutations in C1 inhibitor (C1INH, *SERPING1*). In type I HAE, found in 85% of patients, plasma levels of C1INH are less than 35% of normal, leading to a loss of function of the C1INH [46–48]. In HAE type II, the C1INH serum levels are normal or elevated, but the protein is non-functional due to a mutation within the RCL domain that also causes the inefficient inhibition of the target protease [48]. Antithrombin deficiency (AT3D, #MIM613118), which is the result of variants of the inhibitory serpin antithrombin III (AT3, *SERPINC1*), leads to venous thromboembolic disease. As for the HAE, two categories of AT3D have been defined based on AT3 levels in the plasma and inhibitory activity [49]. Most AT3D manifestation belong to the type I deficiency group with a severe plasma deficiency of AT3; in type II (functional) deficiency, these subjects possess normal serum level for AT3, but mutations in functional domains of this anticoagulant, including the RCL, the heparin-binding site (HBS), or the A- and C-sheet domain, impair or abolish inhibitory activity [50–53].

We now report AAT Iners as the first-described, pure type II AATD variant. We have shown that this variant is secreted at wild-type levels and would therefore not be identified in a patient by conventionally used diagnostic protocols. However, as it is non-functional, a carrier is likely to present the same susceptibility to lung disease as individuals with a recognised deficiency mutant. AATD type II mutants are likely to contribute to the under-diagnosed burden of disease in the general population.

## Methods

*Reagents and antibodies.* Product details are listed in S1 Table.

### Identification of G349R in population databases

The nucleotide (GGG>AGG) encoding the G349R AAT variant (rs12077) was identified in the free-access exome database ExAC v.0.3.1 (60,706 subjects) [54] (http://exac.broadinstitute.org/) and in dbSNP [55].

### Multiple sequence alignments

Multiple sequence alignments were performed with Clustal Omega (https://www.ebi.ac.uk/Tools/msa/clustalo/) using the protein sequence of human alpha-1-antitrypsin (Uniprot P01009), SERPINA1 orthologues and other human serpins (for the complete list of the results see S1 File). Conservation patterns were generated with WebLogo Server V3.6.0 [56].

#### Expression vectors

The mammalian expression vectors for expression of AAT variants are based on pcDNA3.1/Zeo (+) [29]. Bacterial expression of hexahistidine-tagged protein was undertaken using pQE-30 (Qiagen). The AAT Iners mutation was obtained using the QuikChange II Site-Directed Mutagenesis Kit and the oligonucleotide 5’-taaaaacatggccctagcagcttcagtccctttct (and reverse complement thereof).

### Bacterial expression and recombinant protein purification

Recombinant proteins were expressed in the XL1-Blue strain of *E. coli* and purified by nickel-affinity chromatography and ion-exchange chromatography as described previously [38]. Purity was assessed using 4-12% w/v acrylamide SDS-PAGE and 3-12% w/v acrylamide non-denaturing PAGE (Life Technologies) and the resulting proteins were exchanged into 20 mM Tris-HCl pH 7.4, 100 mM NaCl and stored at −80°C.

### Cell culture and transfection

HEK 293T/17 (ATCC, CRL-11268) and Hepa 1-6 (ATCC, CRL-1830) cells were maintained in DMEM/10% v/v FBS. Transfections with vectors encoding M1V, Z or AAT Iners were performed with PEI “Max” or with FuGENE HD as described previously [31,57,58]. To analyse AAT in the cell media, transfected cells were incubated in serum-free Optimem for 24h at 37°C. Cell media were collected and centrifuged at 800g for 5’. Soluble and insoluble cellular fractions were obtained by lysing cells in 10 mM Tris-HCl pH 7.4, 150 mM NaCl, 1% v/v NP-40 and a protease inhibitor cocktail (BLA) with subsequent centrifugation at 16000g to separate the soluble to the insoluble fraction.

### SDS-PAGE, non-denaturating PAGE and immunoblot

Cell media and intracellular fractions were resolved by 7.5% w/v acrylamide SDS-PAGE or by 8% w/v acrylamide non-denaturing as previously described [30,32]. The gels were blotted to PVDF 0. 45 μm membranes by wet transfer, probed with the indicated primary antibodies, revealed with HRP-conjugated secondary antibodies and detected by ECL Clarity and exposure to Hyperfilm ECL.

### Sandwich ELISA

Quantification of AAT in cell media was performed by sandwich ELISA as previously described [32], using rabbit anti-AAT polyclonal antibody (pAb) for capture and HRP-conjugated sheep anti-AAT pAb for detection. AAT concentrations were calculated for each experiment using a standard curve of commercial purified AAT (Millipore) and expressed as percentages of M AAT concentration.

### Formation of the inhibitory complex between AAT and serine-proteases

Culture media of HEK293T cells containing 10 ng of M1V or AAT Iners variants were incubated at 37°C for 20 min with 1:1 and 1:2 molar ratios of A1AT:HNE in 10 mM phosphate buffer pH 7.4/50 mM NaCl, before separation on 7.5% w/v acrylamide SDS-PAGE and immunoblot with anti-A1AT pAb. Recombinant wild-type M1V and AAT Iners produced in bacteria were incubated at 37°C for 30 min with equimolar purified human plasma kallikrein (KLK) in a buffer containing 10 mM Tris-HCl pH 8.0, 50 mM NaCl, 0.02% w/v PEG8000. Samples were then separated by 4-12% w/v acrylamide SDS-PAGE and revealed with Coomassie brilliant blue.

### Assessment of proteinase inhibition

The stoichiometry of inhibition (SI) was assessed at 25°C using bovine α-chymotrypsin or HNE as described previously [42].

### Molecular dynamics analysis

Molecular dynamics (MD) simulations of the wild-type and G349R mutant protein were performed using the GROMACS software package [59,60]. The initial structure for the wild-type protein was taken from PDB 1QLP. The mutant structure was built from the wild-type, by replacing residue 349 with an arginine. The force field amber99-sb was used, with the PME method for Coulomb interactions and a Lennard-Jones potential with a cut-off of 10 Å for the short-range interactions. The initial structure was completed by addition of hydrogens and solvated with TIP3P water in a simulation box with a minimum distance of 10 Å between solute and box boundaries. Na+ and Cl^−^ ions were added to reproduce a salt concentration of 150 mM and to neutralize the system. All simulations were conducted at 310 K. The system was first subjected to energy minimization, then equilibration at constant volume for 100 ps, and at constant pressure for 100 ps was performed before the production run at constant temperature and pressure. The temperature was kept constant by velocity rescaling with a characteristic time of 0.1 ps. The pressure was controlled using the Parrinello-Rahman method with a time constant of 1 ps and a compressibility of 4.5-10^−5^ bar^−1^.

### Statistical analysis

All the statistical analyses were performed by software Prism5 (GraphPad software Inc, San Diego, USA) as detailed in the figure legends.

## Abbreviations

AAT: alpha-1-antitrypsin
AATD: alpha-1-antitrypsin deficiency
HNE: neutrophils elastase
RCL: Reactive Centre Loop
MD: Molecular Dynamics
pAb: polyclonal antibody

## Author contributions

Investigation: ML, EE, FG, RB, JAI, AF

Writing: ML, FG, DAL, JAI, AF

## Acknowledgements

The authors thank Dr. Edoardo Giacopuzzi (University of Brescia, Italy) and Dr. Ilaria Ferrarotti (University of Pavia, Italy) for discussions; we also acknowledge Leonardo Lanfranchi and Giovanni Sgrò (University of Brescia, Italy) for technical assistance.

## Supporting information

**S1 Video:** Movie showing one nanosecond of an MD trajectory of G349R AAT, during which the side chain of M358 changes orientation.

Residues M358 and R349 are highlighted in ball-and-stick representation, while the secondary structure is shown for the rest of the protein. The movie was generated by means of VMD (http://www.ks.uiuc.edu/Research/vmd/).

**S1 Table:** Reagents and antibodies.

**S1 File:** Alignments by Clustal Omega of the human alpha-1-antitrypsin (Uniprot P01009) with SERPINA1 orthologues or inhibitory and non-inhibitory human serpins.

## References

1. Irving JA, Pike RN, Lesk AM, Whisstock JC. Phylogeny of the serpin superfamily: implications of patterns of amino acid conservation for structure and function. Genome Res. 2000;10: 1845–1864.

2. Beatty K, Bieth J, Travis J. Kinetics of association of serine proteinases with native and oxidized alpha-1-proteinase inhibitor and alpha-1-antichymotrypsin. J Biol Chem. 1980;255: 3931–3934.

3. Rao NV, Wehner NG, Marshall BC, Gray WR, Gray BH, Hoidal JR. Characterization of proteinase-3 (PR-3), a neutrophil serine proteinase. Structural and functional properties. J Biol Chem. 1991;266: 9540–9548.

4. Crystal RG. Alpha 1-antitrypsin deficiency, emphysema, and liver disease. Genetic basis and strategies for therapy. J Clin Invest. 1990;85: 1343–1352. doi:10.1172/JCI114578

5. Lomas DA, Evans DL, Finch JT, Carrell RW. The mechanism of Z alpha 1-antitrypsin accumulation in the liver. Nature. 1992;357: 605–607. doi:10.1038/357605a0

6. American Thoracic Society, European Respiratory Society. American Thoracic Society/European Respiratory Society statement: standards for the diagnosis and management of individuals with alpha-1 antitrypsin deficiency. Am J Respir Crit Care Med. 2003;168: 818–900. doi:10.1164/rccm.168.7.818

7. Fra A, Cosmi F, Ordoñez A, Berardelli R, Perez J, Guadagno NA, et al. Polymers of Z α1-antitrypsin are secreted in cell models of disease. Eur Respir J. 2016;47: 1005–1009. doi:10.1183/13993003.00940–2015

8. Tan L, Dickens JA, Demeo DL, Miranda E, Perez J, Rashid ST, et al. Circulating polymers in α1-antitrypsin deficiency. Eur Respir J. 2014;43: 1501–1504. doi:10.1183/09031936.00111213

9. Mahadeva R, Atkinson C, Li Z, Stewart S, Janciauskiene S, Kelley DG, et al. Polymers of Z alpha1-antitrypsin co-localize with neutrophils in emphysematous alveoli and are chemotactic in vivo. Am J Pathol. 2005;166: 377–386.

10. Schechter I, Berger A. On the size of the active site in proteases. I. Papain. 1967. Biochem Biophys Res Commun. 2012;425: 497–502. doi:10.1016/j.bbrc.2012.08.015

11. Dementiev A, Dobó J, Gettins PGW. Active site distortion is sufficient for proteinase inhibition by serpins: structure of the covalent complex of alpha1-proteinase inhibitor with porcine pancreatic elastase. J Biol Chem. 2006;281: 3452–3457. doi:10.1074/jbc.M510564200

12. Strickland DK, Ashcom JD, Williams S, Burgess WH, Migliorini M, Argraves WS. Sequence identity between the alpha 2-macroglobulin receptor and low density lipoprotein receptor-related protein suggests that this molecule is a multifunctional receptor. J Biol Chem. 1990;265: 17401–17404.

13. Huntington JA, Read RJ, Carrell RW. Structure of a serpin-protease complex shows inhibition by deformation. Nature. 2000;407: 923–926. doi:10.1038/35038119

14. Andersen OJ, Risør MW, Poulsen EC, Nielsen NC, Miao Y, Enghild JJ, et al. Reactive Center Loop Insertion in α-l-Antitrypsin Captured by Accelerated Molecular Dynamics Simulation. Biochemistry. 2017;56: 634–646. doi:10.1021/acs.biochem.6b00839

15. Maddur AA, Swanson R, Izaguirre G, Gettins PGW, Olson ST. Kinetic intermediates en route to the final serpin-protease complex: studies of complexes of α1-protease inhibitor with trypsin. J Biol Chem. 2013;288: 32020–32035. doi:10.1074/jbc.M113.510990

16. Stratikos E, Gettins PG. Formation of the covalent serpin-proteinase complex involves translocation of the proteinase by more than 70 A and full insertion of the reactive center loop into beta-sheet A. Proc Natl Acad Sci U S A. 1999;96: 4808–4813.

17. Ye S, Cech AL, Belmares R, Bergstrom RC, Tong Y, Corey DR, et al. The structure of a Michaelis serpin-protease complex. Nat Struct Biol. 2001;8: 979–983. doi:10.1038/nsb1101-979

18. Peterson FC, Gordon NC, Gettins PG. Formation of a noncovalent serpin-proteinase complex involves no conformational change in the serpin. Use of 1H-15N HSQC NMR as a sensitive nonperturbing monitor of conformation. Biochemistry. 2000;39: 11884–11892.

19. Lawrence DA, Olson ST, Muhammad S, Day DE, Kvassman JO, Ginsburg D, et al. Partitioning of serpin-proteinase reactions between stable inhibition and substrate cleavage is regulated by the rate of serpin reactive center loop insertion into beta-sheet A. J Biol Chem. 2000;275: 5839–5844.

20. Lomas DA, Evans DL, Stone SR, Chang WS, Carrell RW. Effect of the Z mutation on the physical and inhibitory properties of alpha 1-antitrypsin. Biochemistry. 1993;32: 500–508.

21. Cook L, Burdon JG, Brenton S, Knight KR, Janus ED. Kinetic characterisation of alpha-1-antitrypsin F as an inhibitor of human neutrophil elastase. Pathology (Phila). 1996;28: 242–247.

22. Okayama H, Brantly M, Holmes M, Crystal RG. Characterization of the molecular basis of the alpha 1-antitrypsin F allele. Am J Hum Genet. 1991;48: 1154–1158.

23. Nyon MP, Segu L, Cabrita LD, Lévy GR, Kirkpatrick J, Roussel BD, et al. Structural dynamics associated with intermediate formation in an archetypal conformational disease. Struct Lond Engl 1993. 2012;20: 504–512. doi:10.1016/j.str.2012.01.012

24. Haq I, Irving JA, Saleh AD, Dron L, Regan-Mochrie GL, Motamedi-Shad N, et al. Deficiency Mutations of Alpha-1 Antitrypsin. Effects on Folding, Function, and Polymerization. Am J Respir Cell Mol Biol. 2016;54: 71–80. doi:10.1165/rcmb.2015-0154OC

25. Owen MC, Brennan SO, Lewis JH, Carrell RW. Mutation of antitrypsin to antithrombin. alpha 1-antitrypsin Pittsburgh (358 Met leads to Arg), a fatal bleeding disorder. N Engl J Med. 1983;309: 694–698. doi:10.1056/NEJM198309223091203

26. Karczewski KJ, Weisburd B, Thomas B, Solomonson M, Ruderfer DM, Kavanagh D, et al. The ExAC browser: displaying reference data information from over 60 000 exomes. Nucleic Acids Res. 2017;45: D840–D845. doi:10.1093/nar/gkw971

27. Giacopuzzi E, Laffranchi M, Berardelli R, Ravasio V, Ferrarotti I, Gooptu B, et al. Real-world clinical applicability of pathogenicity predictors assessed on SERPINA1 mutations in alpha-1-antitrypsin deficiency. Hum Mutat. 2018;39: 1203–1213. doi:10.1002/humu.23562

28. Hopkins PC, Carrell RW, Stone SR. Effects of mutations in the hinge region of serpins. Biochemistry. 1993;32: 7650–7657.

29. Medicina D, Montani N, Fra AM, Tiberio L, Corda L, Miranda E, et al. Molecular characterization of the new defective P(brescia) alpha1-antitrypsin allele. Hum Mutat. 2009;30: E771–781. doi:10.1002/humu.21043

30. Fra AM, Gooptu B, Ferrarotti I, Miranda E, Scabini R, Ronzoni R, et al. Three new alpha1-antitrypsin deficiency variants help to define a C-terminal region regulating conformational change and polymerization. PloS One. 2012;7: e38405. doi:10.1371/journal.pone.0038405

31. Ronzoni R, Berardelli R, Medicina D, Sitia R, Gooptu B, Fra AM. Aberrant disulphide bonding contributes to the ER retention of alpha1-antitrypsin deficiency variants. Hum Mol Genet. 2016;25: 642–650. doi:10.1093/hmg/ddv501

32. Miranda E, Ferrarotti I, Berardelli R, Laffranchi M, Cerea M, Gangemi F, et al. The pathological Trento variant of alpha-1-antitrypsin (E75V) shows nonclassical behaviour during polymerization. FEBS J. 2017;284: 2110–2126. doi:10.1111/febs.14111

33. Laffranchi M, Berardelli R, Ronzoni R, Lomas DA, Fra A. Hetero-polymerization of α1-antitrypsin mutants in cell models mimicking heterozygosity. Hum Mol Genet. 2018; doi:10.1093/hmg/ddy090

34. Teckman JH, Perlmutter DH. The endoplasmic reticulum degradation pathway for mutant secretory proteins alpha1-antitrypsin Z and S is distinct from that for an unassembled membrane protein. J Biol Chem. 1996;271: 13215–13220.

35. Miranda E, Römisch K, Lomas DA. Mutants of neuroserpin that cause dementia accumulate as polymers within the endoplasmic reticulum. J Biol Chem. 2004;279: 28283–28291. doi:10.1074/jbc.M313166200

36. Lukacs CM, Rubin H, Christianson DW. Engineering an Anion-Binding Cavity in Antichymotrypsin Modulates the “Spring-Loaded” Serpin-Protease Interaction. Biochemistry. 1998;37: 3297–3304. doi:10.1021/bi972359e

37. Rawlings ND, Waller M, Barrett AJ, Bateman A. MEROPS: the database of proteolytic enzymes, their substrates and inhibitors. Nucleic Acids Res. 2014;42: D503–509. doi:10.1093/nar/gkt953

38. Zhou A, Carrell RW, Huntington JA. The serpin inhibitory mechanism is critically dependent on the length of the reactive center loop. J Biol Chem. 2001;276: 27541–27547. doi:10.1074/jbc.M102594200

39. Fokkema IFAC, Taschner PEM, Schaafsma GCP, Celli J, Laros JFJ, den Dunnen JT. LOVD v.2.0: the next generation in gene variant databases. Hum Mutat. 2011;32: 557–563. doi:10.1002/humu.21438

40. Huntington JA, Fan B, Karlsson KE, Deinum J, Lawrence DA, Gettins PG. Serpin conformational change in ovalbumin. Enhanced reactive center loop insertion through hinge region mutations. Biochemistry. 1997;36: 5432–5440. doi:10.1021/bi9702142

41. Yamasaki M, Arii Y, Mikami B, Hirose M. Loop-inserted and thermostabilized structure of P1-P1’ cleaved ovalbumin mutant R339T. J Mol Biol. 2002;315: 113–120. doi:10.1006/jmbi.2001.5056

42. Haq I, Irving JA, Faull SV, Dickens JA, Ordóñez A, Belorgey D, et al. Reactive centre loop mutants of α-1-antitrypsin reveal position-specific effects on intermediate formation along the polymerization pathway. Biosci Rep. 2013;33. doi:10.1042/BSR20130038

43. Yamasaki M, Sendall TJ, Harris LE, Lewis GMW, Huntington JA. Loop-sheet mechanism of serpin polymerization tested by reactive center loop mutations. J Biol Chem. 2010;285: 30752–30758. doi:10.1074/jbc.M110.156042

44. Bhoola KD, Figueroa CD, Worthy K. Bioregulation of kinins: kallikreins, kininogens, and kininases. Pharmacol Rev. 1992;44: 1–80.

45. Jörg M, Binder BR. Kinetic analysis of plasminogen activation by purified plasma kallikrein. Thromb Res. 1985;39: 323–331.

46. Carugati A, Pappalardo E, Zingale LC, Cicardi M. C1-inhibitor deficiency and angioedema. Mol Immunol. 2001;38: 161–173.

47. Caccia S, Suffritti C, Carzaniga T, Berardelli R, Berra S, Martorana V, et al. Intermittent C1-Inhibitor Deficiency Associated with Recessive Inheritance: Functional and Structural Insight. Sci Rep. 2018;8: 977. doi:10.1038/s41598-017-16667-w

48. Pappalardo E, Caccia S, Suffritti C, Tordai A, Zingale LC, Cicardi M. Mutation screening of C1 inhibitor gene in 108 unrelated families with hereditary angioedema: functional and structural correlates. Mol Immunol. 2008;45: 3536–3544. doi:10.1016/j.molimm.2008.05.007

49. Corral J, de la Morena-Barrio ME, Vicente V. The genetics of antithrombin. Thromb Res. 2018;169: 23–29. doi:10.1016/j.thromres.2018.07.008

50. Águila S, Izaguirre G, Martínez-Martínez I, Vicente V, Olson ST, Corral J. Disease-causing mutations in the serpin antithrombin reveal a key domain critical for inhibiting protease activities. J Biol Chem. 2017;292: 16513–16520. doi:10.1074/jbc.M117.787325

51. Chuang YJ, Gettins PG, Olson ST. Importance of the P2 glycine of antithrombin in target proteinase specificity, heparin activation, and the efficiency of proteinase trapping as revealed by a P2 Gly --> Pro mutation. J Biol Chem. 1999;274: 28142–28149.

52. Lane DA, Olds RJ, Conard J, Boisclair M, Bock SC, Hultin M, et al. Pleiotropic effects of antithrombin strand 1C substitution mutations. J Clin Invest. 1992;90: 2422–2433. doi:10.1172/JCI116133

53. Koide T, Odani S, Takahashi K, Ono T, Sakuragawa N. Antithrombin III Toyama: replacement of arginine-47 by cysteine in hereditary abnormal antithrombin III that lacks heparin-binding ability. Proc Natl Acad Sci U S A. 1984;81: 289–293.

54. Lek M, Karczewski KJ, Minikel EV, Samocha KE, Banks E, Fennell T, et al. Analysis of protein-coding genetic variation in 60,706 humans. Nature. 2016;536: 285–291. doi:10.1038/nature19057

55. Sherry ST, Ward MH, Kholodov M, Baker J, Phan L, Smigielski EM, et al. dbSNP: the NCBI database of genetic variation. Nucleic Acids Res. 2001;29: 308–311.

56. Crooks GE, Hon G, Chandonia J-M, Brenner SE. WebLogo: a sequence logo generator. Genome Res. 2004;14: 1188–1190. doi:10.1101/gr.849004

57. Fra A, D’Acunto E, Laffranchi M, Miranda E. Cellular Models for the Serpinopathies. Methods Mol Biol Clifton NJ. 2018;1826: 109–121. doi:10.1007/978-1-4939-8645-3_7

58. Tiberio L, Nascimbeni R, Villanacci V, Casella C, Fra A, Vezzoli V, et al. The decrease of mineralcorticoid receptor drives angiogenic pathways in colorectal cancer. PloS One. 2013;8: e59410. doi:10.1371/journal.pone.0059410

59. Hess B, Kutzner C, van der Spoel D, Lindahl E. GROMACS 4: Algorithms for Highly Efficient, Load-Balanced, and Scalable Molecular Simulation. J Chem Theory Comput. 2008;4: 435–447. doi:10.1021/ct700301q

60. Abraham MJ, Murtola T, Schulz R, Páll S, Smith JC, Hess B, et al. GROMACS: High performance molecular simulations through multi-level parallelism from laptops to supercomputers. SoftwareX. 2015;1–2: 19–25. doi:10.1016/j.softx.2015.06.001

